# A signature of neural coding at human perceptual limits

**DOI:** 10.1101/051714

**Authors:** Paul M Bays

## Abstract

Simple visual features, such as orientation, are thought to be represented in the spiking of visual neurons using population codes. I show that optimal decoding of such activity predicts characteristic deviations from the normal distribution of errors at low gains. Examining human perception of orientation stimuli, I show that these predicted deviations are present at near-threshold levels of contrast. The findings may provide a neural-level explanation for the appearance of a threshold in perceptual awareness, whereby stimuli are categorized as seen or unseen. As well as varying in error magnitude, perceptual judgments differ in certainty about what was observed. I demonstrate that variations in the total spiking activity of a neural population can account for the empirical relationship between subjective confidence and precision. These results establish population coding and decoding as the neural basis of perception and perceptual confidence.

## Introduction

Population coding describes a method by which information can be encoded in, and recovered from, the combined activity of a pool of neurons (Georgopoulos et al., 1982; Pouget, Dayan, & Zemel, 2000; Salinas & Abbott, 1994; Seung & Sompolinsky, 1993; Vogels, 1990). For example, in area V1, simple cells’ spiking activity contains information about the orientation of visual stimuli. Each neuron’s mean firing rate is described by an approximately bell-shaped tuning curve, with a maximum at the cell’s ‘preferred’ orientation. This orientation varies from neuron to neuron, and the population as a whole encodes information about every possible orientation. Simple neural mechanisms have been proposed that can decode population spiking activity and recover the information about the stimulus (Deneve, Latham, & Pouget, 1999; Jazayeri & Movshon, 2006), although these theoretical mechanisms have not as yet been validated by neurophysiology. Irrespective of mechanism, the decoded values are necessarily noisy approximations to the stimulus, due to the stochastic nature of spiking events.

The principle that internal noise is responsible for errors in detection or discrimination of visual patterns has a long history in vision science (e.g. Pelli, 1985), and many models have been proposed to account for behavioral performance on such tasks, incorporating varying degrees of biological detail from simple linear filters to spiking neurons (e.g. Bradley, Abrams, & Geisler, 2014; Foley et al., 2007; Goris et al., 2013; Itti, Koch, & Braun, 2000; Watson & Ahumada, 2005). The present study diverges from previous work by examining predictions of a population coding model for the distribution of errors in an estimation task. Variability in perception of visual stimuli is typically assumed to follow a normal distribution (Green & Swets, 1966; Swets, Tanner Jr, & Birdsall, 1961); the normal is a central limit distribution, a distribution to which values converge when many small influences are summed together, and for this reason it is ubiquitous in biology. However, here I show that the mathematics of population coding puts it in conflict with the assumption of normality. Specifically, characteristic deviations from the normal distribution are predicted at low gains, i.e. when spiking activity is reduced. I confirm the presence of these deviations in human estimation of low contrast stimuli, demonstrating a causal connection between population coding and perception.

As well as explaining errors, the neural model predicts variation in the certainty associated with each judgment, i.e. some estimates are more reliable than others. I show that observers have access to reliability information and use it to assign confidence to their perceptions. Previous attempts to explain the accuracy of confidence judgments have proposed a relationship to response time (Audley, 1960), or to the balance of accumulated evidence favoring one response over another (Smith & Vickers, 1988; Vickers & Packer, 1982). Here I show that the sum of spiking activity in the population encoding a stimulus could provide a plausible neural basis for confidence judgments.

## Experimental Procedures

### Experiment

Eight participants (1 male, 7 females; aged 22–41 years) participated in the study after giving informed consent, in accordance with the Declaration of Helsinki. All participants reported normal color vision and had normal or corrected-to-normal visual acuity. Stimuli were presented on a 21-inch linearized CRT monitor with a refresh rate of 130 Hz. The monitor was fitted with a neutral density filter to decrease the luminance range to the level of human detection thresholds. Participants sat with their head supported by a forehead and chin rest and viewed the monitor at a distance of 60 cm.

Stimuli consisted of Gabor patches of varying contrast and orientation (wavelength of sinusoid, 0.75° of visual angle; s.d. of Gaussian envelope, 0.75°), presented at display center on a gray background. Stimuli were presented within an annulus (white, radius 4°) which was always present on the display.

Detection thresholds were obtained prior to the main experiment using an adaptive estimation method. On each trial (160 in total) a Gabor was presented for 100 ms randomly at one of two time-points, 1 s apart, identified by auditory cues; participants reported at which of the two timepoints the Gabor was present. Detection threshold was defined as the Gabor contrast at which participants performed at 75% correct, estimated by fitting a sigmoid function to the contrast-response data. Gabor contrast was selected on each trial to maximize the information available for this estimation (Psi method; Kontsevich & Tyler, 1999).

In the main experiment, each trial began with presentation of a randomly-oriented Gabor patch for 100 ms and a simultaneous auditory tone. The contrast of the Gabor was chosen at random from {50%, 100%, 200%, 400%} of the previously-obtained detection threshold. After 1 s, a randomly-oriented bar stimulus (white, radius 5°, width 0.1°, central 6° omitted) was overlaid on the annulus; participants adjusted the bar orientation to match the orientation of the Gabor patch, using a computer mouse. They then indicated their confidence in their judgment by clicking on one of a set of buttons labeled {0%, 25%, 50%, 75%, 100%}. Participants completed between 280 and 480 trials.

### Analysis

Orientations were analyzed and are reported with respect to the circular parameter space of possible values, i.e. the space of possible orientations [–90°,90°) was mapped onto the circular space [–π,π) radians. Error for each trial was calculated as the angular deviation between the orientation reported by the participant and the true orientation. Central tendency was assessed using the V statistic for nonuniformity of circular data. Recall precision was defined as 1/*σ*^2^ where 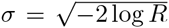 is the circular standard deviation as defined by Fisher (1995), R being the resultant length. Hypotheses regarding the effects of experimental parameters (contrast, subjective confidence rating) were tested with *t* tests.

### Population coding model

I studied encoding and decoding in a population of *M* idealized neurons with orientation tuning and contrast sensitivity. The average response of the *i*th neuron to visual input was defined as (Albrecht & Hamilton, 1982; Carandini & Heeger, 2012; Heeger, 1992):

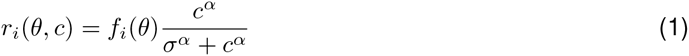

where *θ* is the stimulus orientation, *c* is the stimulus contrast, and *f_i_*(*θ*) is a Von Mises tuning function, centered on *φ_i_*, the neuron’s preferred orientation:

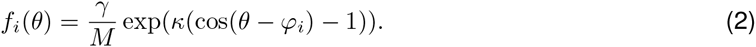

where *γ* is the population gain. Preferred orientations were evenly distributed throughout the range of possible orientations. Spiking activity was modeled as a homogeneous Poisson process, such that the probability of a neuron generating *n* spikes in time *T* was:

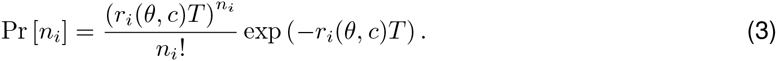

Decoding of orientation information from the population’s spiking activity, n, was based on maximum a posteriori (MAP) decoding. Assuming a uniform prior, this is equivalent to maximizing the likelihood:

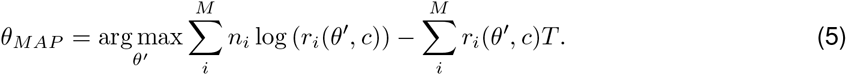

If two or more orientations tied for the maximum, the decoded orientation was sampled at random from the tied values. The output of the model was given by 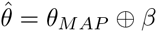, where *β* is a response bias term, and ⊕ indicates addition on the circle. Decoding time *T* was fixed at 100 ms. I considered the limit *M* → ∞. The model therefore has five free parameters: *σ* and *α*, constants of the contrast response function; *γ*, the population gain; *κ*, the tuning curve width; and *β*, the response bias.

#### Fitting the model

While the equations above provide a complete description of the model, further analysis is needed to obtain predictions of the model and fit them to data. From (4),

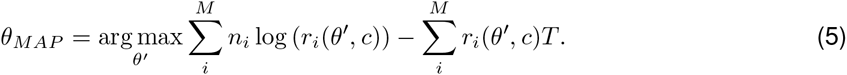

Assuming dense uniform coverage, the second term is constant, so

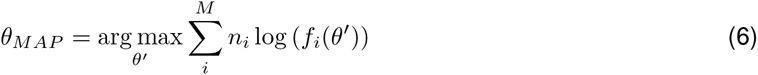

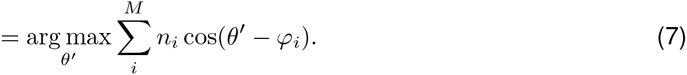

Consider the combined activity of the population in terms of the preferred stimulus corresponding to each spike: {*φ*_(1)_, *φ*_(2)_,… *φ*_(*m*)_} where the notation *φ*_(*i*)_ indicates the preferred orientation of the neuron that generated the *i*th of *m* spikes. The error in the decoded orientation, 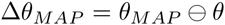, can then be written as,

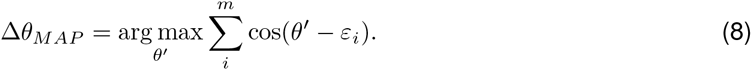

where 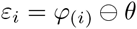. Setting the derivative of the term to be maximized to zero, we obtain,

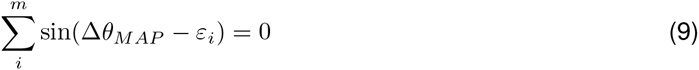

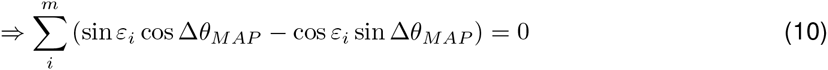

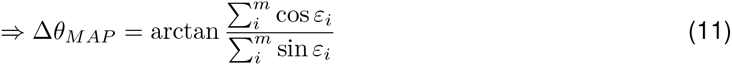

Because spikes are generated by independent Poisson processes, every spike event is conditionally independent of every other given the true stimulus orientation. Approximating the uniformly-spaced discrete distribution of preferred orientations of *M* neurons by a continuous uniform distribution, this probability is given by:

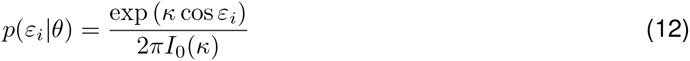

So the error in decoded orientation Δ*θ_MAP_* is the resultant angle (Eq. 11) of a Von Mises (circular normal) random walk (Eq. 12) of *m* steps. It follows that the error for a given resultant length r is Von Mises distributed (Mardia & Jupp, 2009):

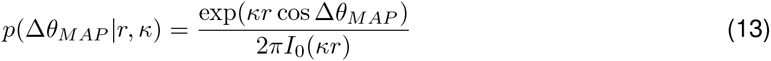

where the distribution of *r* for *m* steps is given by

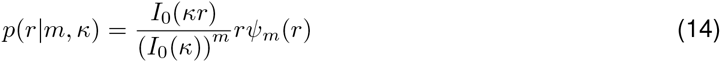

where *rψ_m_*(*r*) is the probability density function for resultant length *r* of a uniform random walk of *m* steps. The distribution of *m*, the total spike count during the decoding interval *T*, being a sum of *M* independent Poisson distributions, is itself Poisson:

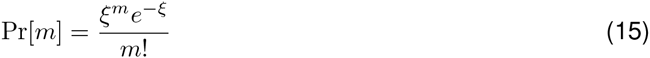

where *ξ* is the expected total spike count,

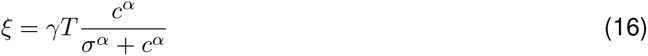

Equations (13), (14) and (15) together provide a means of obtaining the distribution of Δ*θ_MAP_*, and hence of the response error 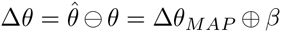, for any values of the free parameters, *σ, α, γ, κ, β*. For *m* ≤ 100, the density *ψ_m_*(*r*) was approximated by Monte Carlo simulation, discretizing over 10^3^ bins. For larger *m* a Gaussian approximation to Equation (14) was used (Mardia & Jupp, 2009):

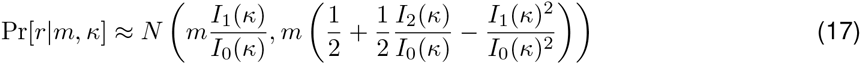

These equations were fit to empirical response data using the Nelder-Mead simplex method (*fminsearch* in MATLAB). Note that, as a mixture of normal distributions of different widths, the distribution of error is in general not normally-distributed.

#### Simulations

To examine predictions of the population coding model in more detail, I performed Monte Carlo simulations (*M* = 100 neurons; 10^5^ repetitions per subject and contrast) using parameters obtained by fitting the model to the experimental data. Note that previous work (Bays, 2014) has shown 100 neurons to be sufficient to approximate the large population limit *M* → ∞: simulating larger numbers of neurons would not have changed the results. Simulated trials were split into two equal bins, according to either the precision of the posterior distribution *p*(*θ*|n) or the total spike count 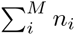, and precision of simulated responses estimated separately for each bin.

#### Modeling detection threshold

I modeled the detection task as follows. On each trial there were two decoding intervals of length 100 ms corresponding to the two timepoints at which a stimulus could be presented. A response was generated according to which interval contained the most spikes. There is no baseline activity in the model neurons so the no-stimulus epoch always contained zero spikes; therefore errors occurred only when the stimulus epoch also had no activity, and then at the guessing rate of 50%. From Eq (15), the probability of generating zero spikes in interval *T* in response to a stimulus of contrast *c* is

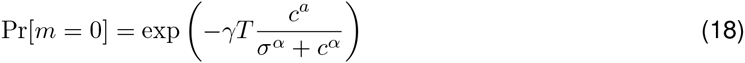

So the threshold contrast at which responses are 75% correct is given by

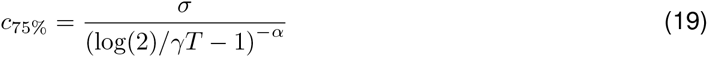

### Threshold model

A threshold model of perceptual judgments would suggest that the stimulus on each trial is either seen, with probability *p*, or not seen, with probability (1 − *p*), where *p* depends on stimulus contrast. Seen stimuli are reported with circular normal (Von Mises) distributed error with standard deviation *σ_seen_* and bias *β*. When the stimulus is not seen the response is random (i.e. drawn from a uniform distribution). The result is a mixture distribution with density:

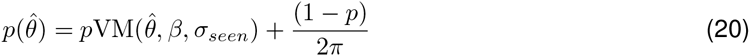

where VM(*θ, μ,σ*) is the Von Mises distribution evaluated at *θ* with mean *μ* and s.d. *σ*. This resulted in a model with six free parameters: *σ_seen_*, *β*, *p*_50%_, *p*_100%_, *p*_200%_, *p*_400%_. Models were compared using AICc (AIC with finite data correction) and BIC.

### Two-stage model

I considered a two-stage model in which the stimulus is first represented with circular normal error before being encoded in the neural population. This could correspond to the case where the nonnormality arises subsequent to initial perceptual representation, for example in working memory. The resulting decoded stimulus estimates are distributed as the convolution of a circular normal with the population coding error distribution obtained above [Equations (13)–(15)]:

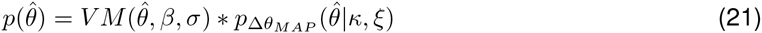

The effect of changing contrast was reflected in the width *σ* of the initial normal representation. This model therefore had seven free parameters: *σ*_50%_, *σ*_100%_, *σ*_200%_, *σ*_400%_, *β, κ*, *ξ*.

### Background activity

I considered a variant of the population coding model in which all neurons have background (baseline) activity, *η*. In this case, the response of the *i*th neuron is given by:

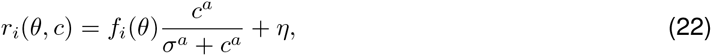

while equations (2)–(4) hold as before. The model of detection is the same as above, except that the no-stimulus epoch contained spikes generated at the baseline rate *η*, while activity in the stimulus epoch was given by Eq (22).

## Results

I examined a model of population coding based on responses of visual cortical neurons to simple oriented stimuli of varying contrast (Fig 1). Mean firing rate of each neuron was determined by the product of its contrast response (Fig 1a), described by a sigmoid relationship between log contrast and firing rate (Albrecht & Hamilton, 1982; Carandini & Heeger, 2012; Heeger, 1992), and its orientation tuning (Fig 1b), described by a bell-shaped tuning function (Pouget, Dayan, & Zemel, 2000). Spikes were generated probabilistically according to a Poisson process (Fig 1c). Estimation of orientation was modeled as maximum a posteriori (MAP) decoding over a fixed temporal window. Because of the noise in spiking activity, the decoded orientation was imprecise with respect to the true stimulus value.

**Figure 1:**
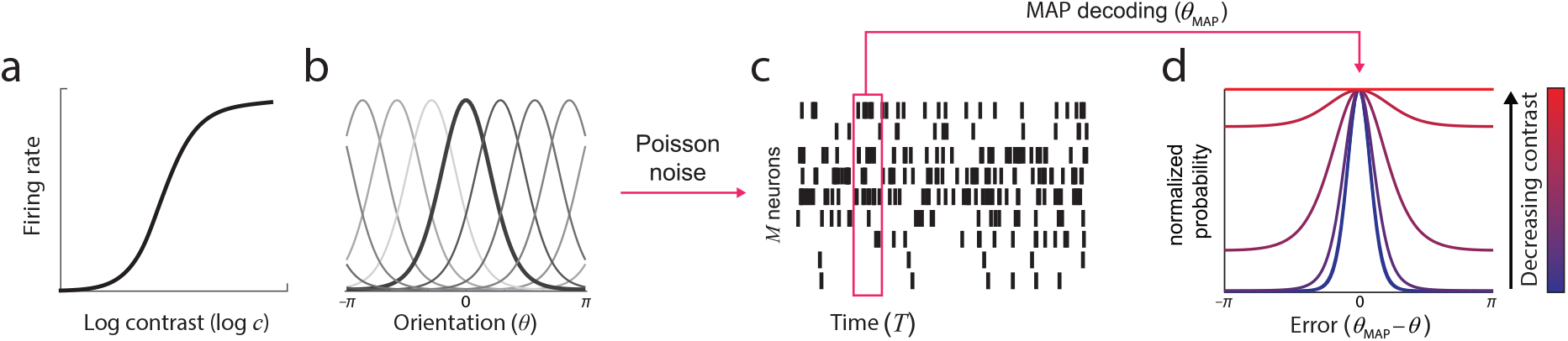
The population coding model. (a & b) Stimulus parameters were encoded in the activity of idealized neurons with contrast sensitivity (a) and bell-shaped orientation tuning functions (b; preferred orientations evenly distributed on the circle). (c) Spikes were generated according to a Poisson process. Estimation of orientation was modeled as MAP decoding of this spiking activity over a fixed time window. (d) Simulations revealed that the distribution of error in the estimated orientation depended on stimulus contrast. At high contrast, errors had an approximately circular normal distribution (e.g. blue curve). As contrast decreased, variability increased and error distributions deviated from circular normality (long tails, e.g. magenta curve). At the lowest contrasts, errors approximated a uniform distribution (e.g. red curve). Error distributions are normalized by peak probability to best illustrate distribution shape.

Modeling results showed that the distribution of errors in the decoded orientation estimate varied with gain, and hence with input contrast (Fig 1d). For high contrast stimuli, the decoded value was distributed approximately as a circular normal (Von Mises) centered on the true orientation (e.g. blue curve). As contrast decreased, the distribution became broader, and also deviated substantially from the circular normal distribution (long tails, e.g. magenta curve). As the contrast fell to zero, the distribution of errors became flatter, approaching the uniform distribution (red line).

### Experimental confirmation

To examine whether non-normality of response errors is a feature of human perceptual judgments, observers were presented with randomly-oriented Gabor patches of varying contrast, at and around each observer’s detection threshold (defined as the contrast at which two-alternative forced choice judgments were 75% correct). They were asked to reproduce the orientation they had seen by rotating a bar stimulus. Figure 2a (black symbols) plots the distribution of response errors for different stimulus contrasts (labeled as percentage of detection threshold). Response precision declined with decreasing contrast, but performance was significantly above chance at every contrast level tested (V > 6.9; t(7) > 2.6, p < 0.032). Significant deviations from circular normality were evident as long tails in the error distribution at detection threshold (100%, highlighted in blue; circular kurtosis of 2.7 greater than circular normal with matched variance; t(7) = 2.8, p = 0.026; also in 8 out of 8 subjects considered individually). Figure 2b plots the discrepancy between the error distributions generated by observers and a circular normal distribution with the same variance.

**Figure 2:**
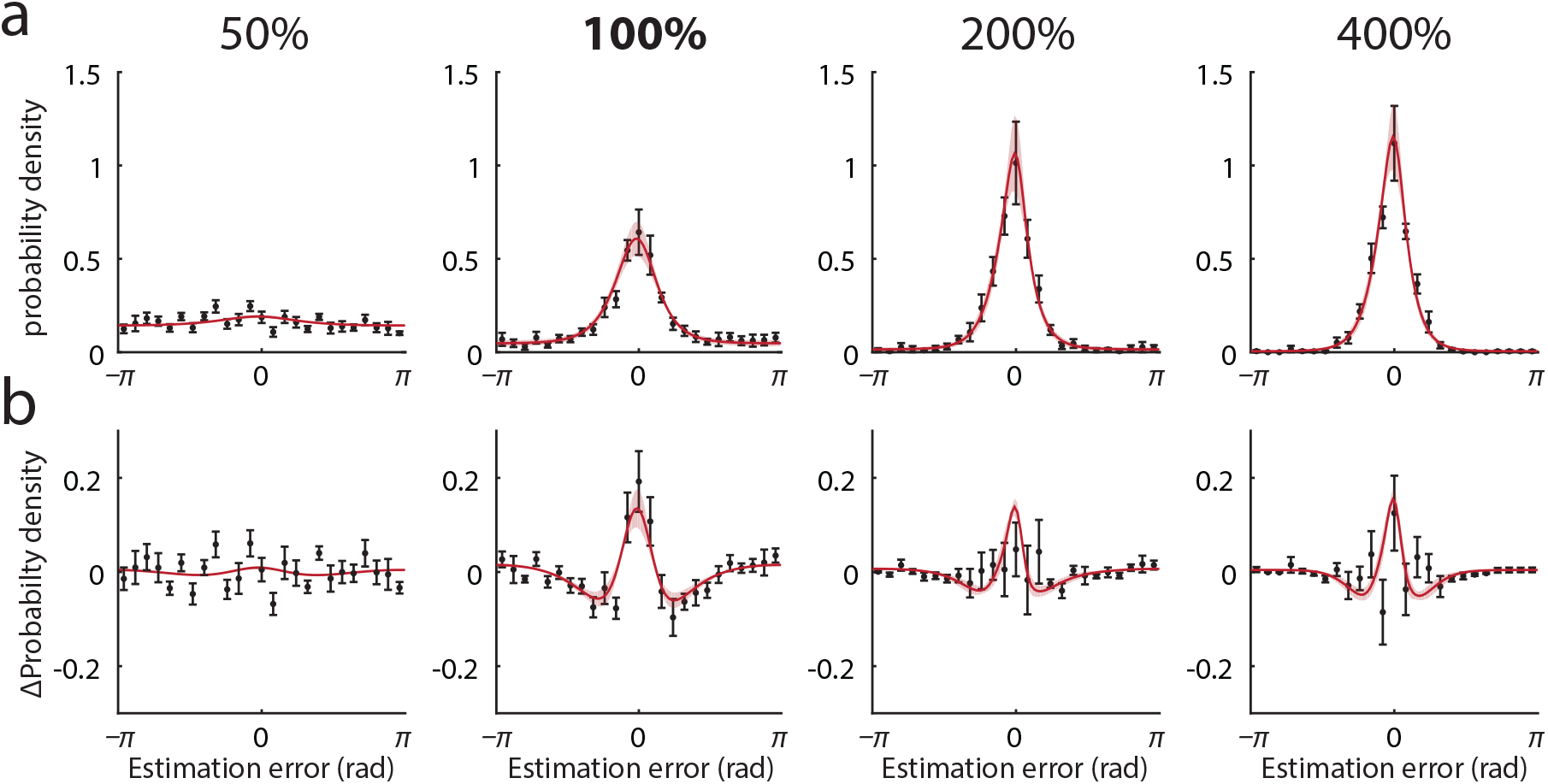
Human estimation errors and model fits. (a) Black symbols show mean distribution of experimental estimation errors (error bars indicate ±1 SE). Different panels correspond to different contrasts, 50%–400% of detection threshold. Note presence of long tails in error distribution at detection threshold (100%). Red curves show mean error distributions for the population coding model with ML parameters (light red patches indicate ±1 SE). (b) Deviation from the circular normal distribution. Black symbols plot mean discrepancy between experimental error frequencies shown in (a) and circular normal (Von Mises) distributions matched in mean and variance. Red curves plot equivalent deviations for the population coding model with ML parameters.

Red curves in Figure 2 show fit of the population coding model (ML parameters: response bias *β* = −0.050 rad ± 0.028 rad, tuning width *κ* = 2.40 ± 0.58, population gain *γ* = 145 Hz ± 92 Hz, contrast response parameters *α* = 48.2 ± 16.6, *σ* = 0.096 ± 0.0081; goodness of fit: r^2^ = 0.64 ± 0.14 SD). The model reproduced both the changes in distribution width with contrast and, importantly, the non-normality of errors around detection threshold. Figure 3 plots response precision as a function of contrast for experimental data (black symbols) and the fitted model (black line).

**Figure 3:**
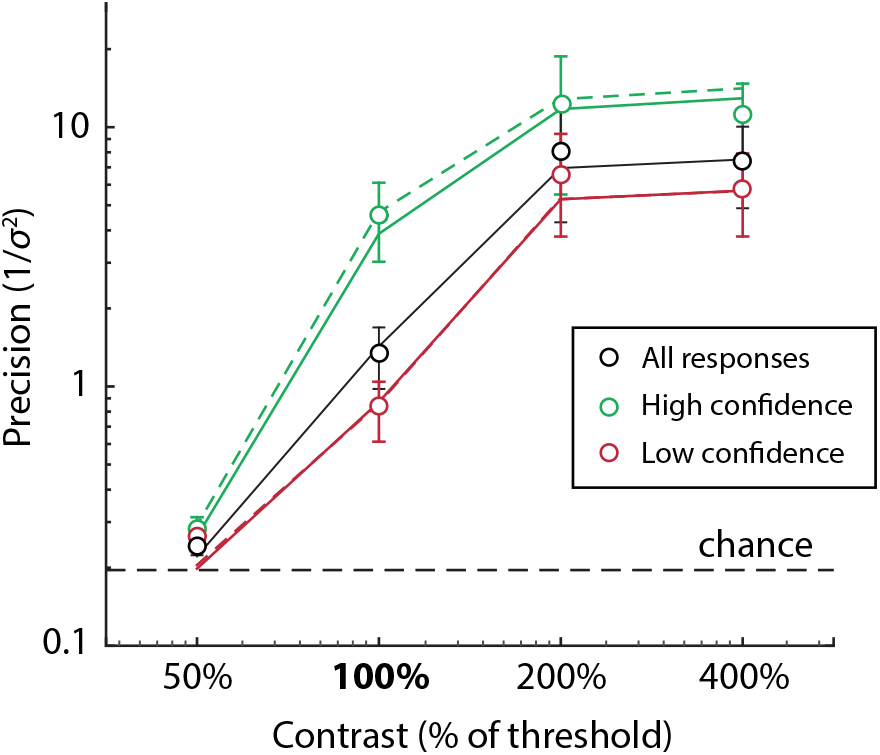
Precision and confidence. Black symbols plot mean precision of experimental responses as a function of contrast (percentage of detection threshold). Dashed black line indicates precision corresponding to random responses. Green and red symbols show mean precision of responses associated with high and low confidence judgments, respectively (median split). Lines show mean precision of simulated responses based on ML parameters. Black line corresponds to all simulated responses, green and red lines correspond to simulations with high and low posterior precision (solid lines) or total spike count (dashed lines), respectively (median split). Note that red and green lines are not fitted to red and green data points.

In addition to perceptual error, the population coding model also makes predictions about stimulus detection. In a two-alternative forced choice task, as used here to estimate detection threshold, a simple observer model selects whichever epoch contained the most spikes. I estimated the threshold contrast that would result in 75% correct responses under this model, based on the ML parameters obtained above. The resulting predictions were statistically indistinguishable from the empirical threshold values (97% ± 16% of empirical threshold contrast, t(7) = 0.19, p = 0.86).

### Other models

I compared the population coding model to a threshold model of perceptual responses (Luce, 1963; Sergent & Dehaene, 2004; Supèr, Spekreijse, & Lamme, 2001), which describes trials as falling into one of two categories: seen and unseen. When the stimulus is seen, responses are distributed normally; when the stimulus is unseen, responses are random. This model generated qualitatively similar predictions to the population coding model, although with a tendency to underestimate non-normality at higher contrasts (Fig 4, blue curves; ML parameters: response bias *β* = −0.044 rad ± 0.027 rad, variability *σ_seen_* = 0.43 ± 0.044, probability seen *p*_50%_ = 0.048 ± 0.016, *p*_100%_ = 0.59 ± 0.11, *p*_200%_ = 0.89 ± 0.08, *p*_400%_ = 0.98 ± 0.013). The threshold model was a poorer fit to the experimental data according to model selection criteria (ΔAICc = 12.6; ΔBIC = 43.5).

**Figure 4:**
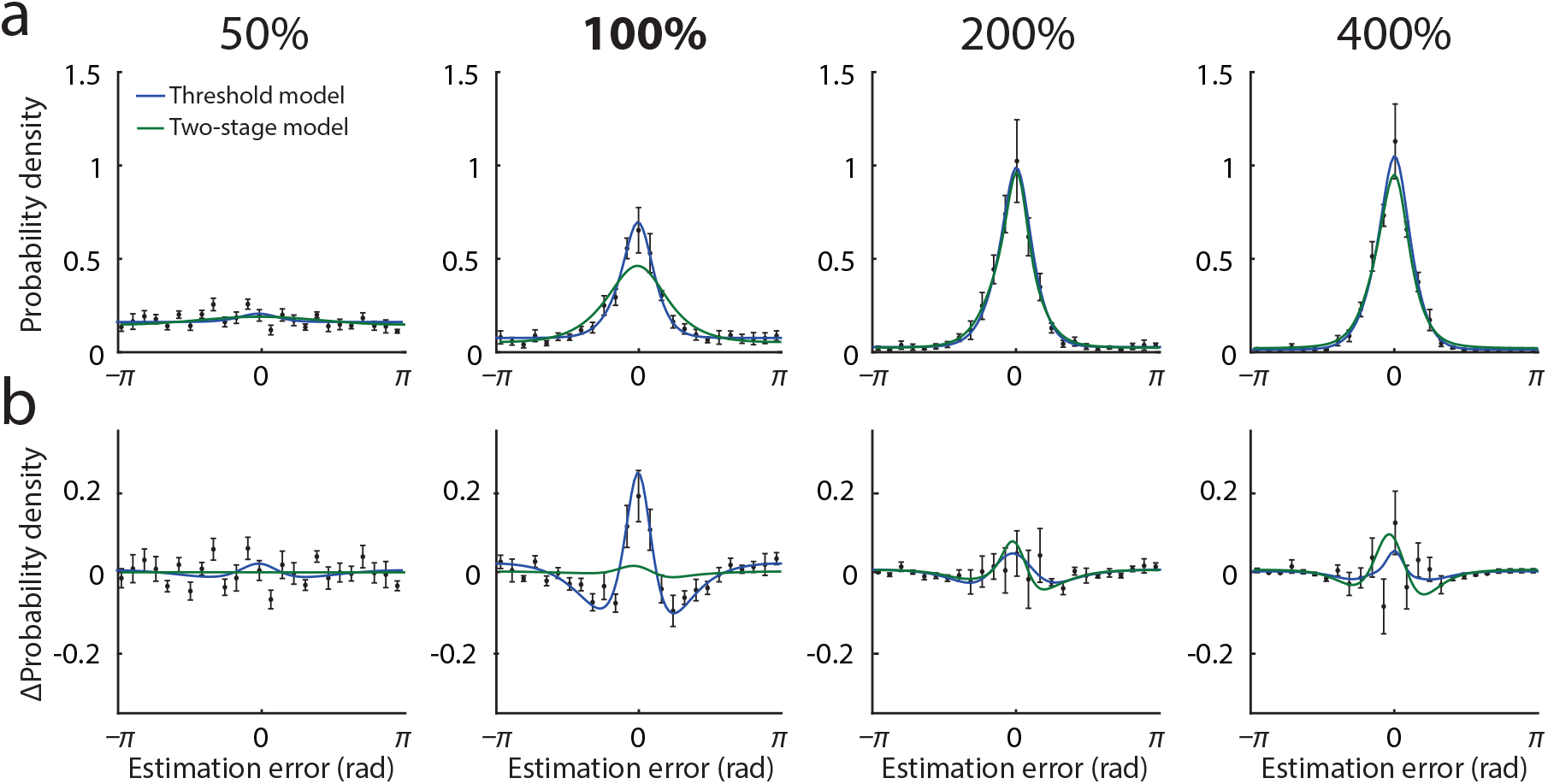
Threshold model. (a) Black symbols show mean distribution of experimental estimation errors (error bars indicate ±1 SE). Different panels correspond to different contrasts, 50%–400% of detection threshold. Blue curves show mean error distributions for the threshold model with ML parameters; green curves show distributions for the two-stage model. (b) Deviation from the circular normal distribution. Black symbols plot mean discrepancy between experimental error frequencies shown in (a) and circular normal (Von Mises) distributions matched in mean and variance. Equivalent deviations are shown for the threshold model (blue) and the two-stage model (green).

Although a standard perceptual task, the orientation reproduction task also has a working memory component, as the target stimulus must be held in mind while the participant adjusts the probe bar. One possibility is that the non-normality arises in working memory storage, subsequent to the perceptual representation. To test this I considered a two-stage model, in which the error arising initially in perception is normally-distributed, with a width determined by the stimulus contrast, and the perceived value is then encoded and decoded according to the population model, introducing non-normality. This model failed to reproduce the non-normality in response distributions, particularly at contrasts around detection threshold (Fig 4, green curves; ML parameters: response bias *β* = −0.062 rad ± 0.023, tuning width *κ* = 17.2 ± 6.1, population activity *ξ* = 13.5 ± 9.4, normal s.d. *σ*_50%_ = 4.0 ± 0.94, *σ*_100%_ = 1.7 ± 0.74, *σ*_200%_ = 0.52 ± 0.14, *σ*_400%_ = 0.48 ± 0.11). The two-stage model was a substantially poorer fit to the experimental data than the population coding model (ΔAICc = 327; ΔBIC = 390).

A final possibility is that non-normality arises from anisotropy in orientation perception. It is well-established that orientation judgements display small biases away from the cardinal angles (e.g. de Gardelle, Kouider, & Sackur, 2010), an ‘anti-Bayesian’ effect possibly due to efficient coding by the underlying neural populations (Wei & Stocker, 2015). As shown in Fig 5a, some evidence for such biases was obtained in the present study, specifically as response shifts away from the horizontal. Because error distributions are calculated by averaging over different stimulus orientations, such biases could potentially result in non-normal distributions of error overall, even if the distribution of error for any given stimulus orientation is normal. To test this, I simulated responses by drawing samples from normal (Von Mises) distributions with the biases and dispersions observed in the data at different stimulus values (15 evenly-spaced bins); Fig 5b (blue curve) plots the resulting deviations from normality in the simulated error distribution. The deviations from normality are an order of magnitude smaller than those observed in the data (black data points), demonstrating that anisotropy in orientation perception cannot account for the non-normality of errors that is the focus of this study.

**Figure 5:**
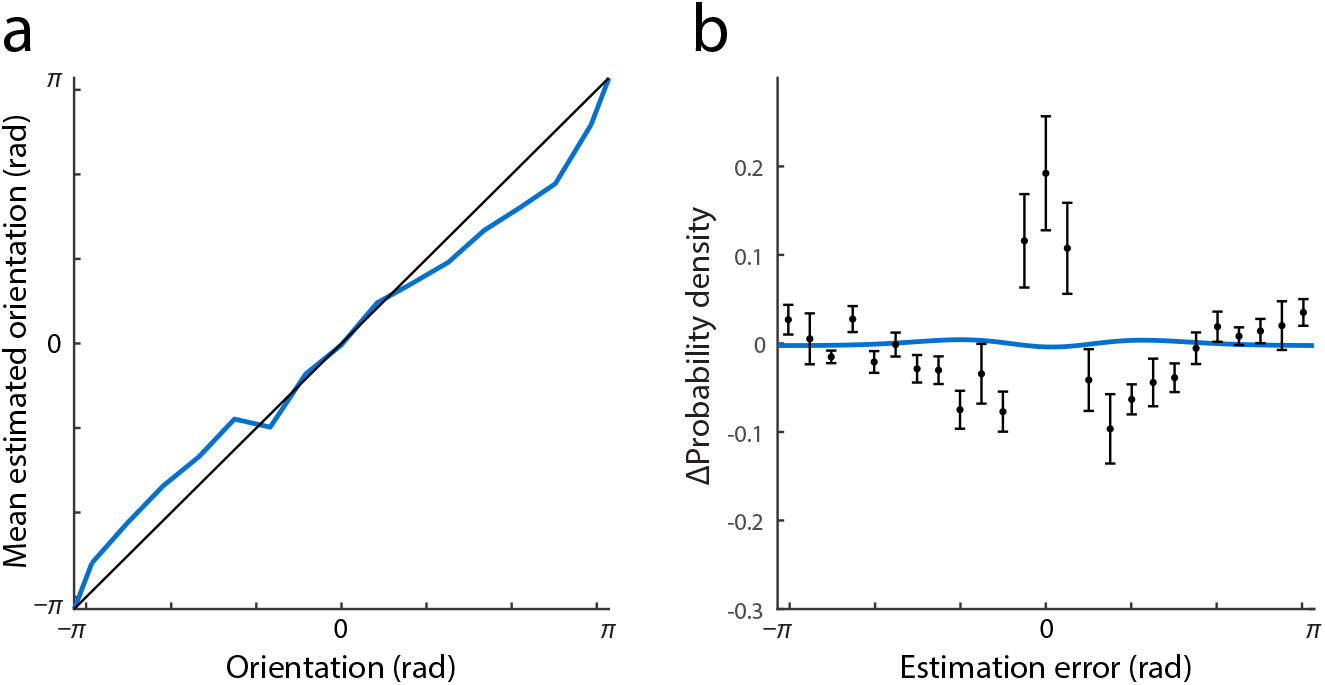
Impact of response biases. (a) Mean estimated orientation at threshold contrast as a function of the true orientation of the stimulus. Black line indicates equality. (b) Blue curve plots estimated deviations from normality resulting from observed orientation biases. Black data points show actual deviations from normality, replotted from Fig 2.

### Subjective confidence

As well as varying in the magnitude of error, responses also varied in the subjective confidence, or reliability, observers assigned to them. Green and red symbols in Figure 3 indicate the precision of high and low confidence responses respectively, based on a median split. Subjective ratings of confidence were significantly correlated with error magnitude for all but the lowest contrast stimuli, indicating that observers had some awareness of the uncertainty in their perception (50% contrast, r^2^ = 0.03, t(7) = 1.0 p = 0.34; 100% contrast, r^2^ = 0.17, t(7) = 4.6, p = 0.003; 200% contrast, r^2^ = 0.07, t(7) = 3.0, p = 0.020; 400% contrast, r^2^ = 0.05, t(7) = 3.6, p = 0.009).

In the population coding model, the parameter that most directly corresponds to response reliability is the precision of the posterior distribution. To assess whether knowledge of this feature of neural decoding could underlie confidence judgments, I performed a median split on the posterior precision of simulated data, generated using the ML parameters obtained above. Green and red solid lines in Figure 3 plot the precision of high and low posterior precision trials respectively. Despite not being fit to the high/low confidence data, this model closely replicated the behavioral results (mean squared error [MSE] 0.073 ± 0.023).

A more directly computable parameter of spiking activity correlated with reliability is the total spike count during the decoding window (Bays, 2014; Ma et al., 2006; Pouget, Dayan, & Zemel, 2003). This parameter was strongly correlated with posterior precision (r^2^ = 0.42). A median split based on total spike count (dashed lines in Fig 3) produced a replication of behavioral results that was indistinguishable from posterior precision (MSE 0.062 ± 0.015, t(7) = 1.3, p = 0.23).

### Background activity

The model of population coding presented above assumes that each neuron’s spiking activity falls to zero at zero contrast. Here I consider the case where all neurons have background (baseline) activity, *η*. This model is considerably less analytically tractable than the no-baseline (*η* = 0) model, and numerically fitting it to the experimental data is impractical. However the predictions of the model share all the main characteristics of the no-baseline case. To illustrate the similarity, I considered the case *η* = 1 Hz. Taking as a starting point the ML parameters of the no-baseline model for a representative observer, I used a grid-search (10 × 10 parameter space, 10^5^ repetitions, *M* = 100) to seek new values of *κ* and *γ* for which the baseline model approximated the predictions of the no-baseline model. As shown in Fig 6, and consistent with previous results (Bays, 2014), the baseline model generated predictions that were almost indistinguishable from those of the no-baseline model, but at higher gain (*γ* = 41.7 Hz, compared to 28.8 Hz in the no-baseline case) and based on broader tuning curves (*κ* = 1.30, compared to 2.12).

**Figure 6:**
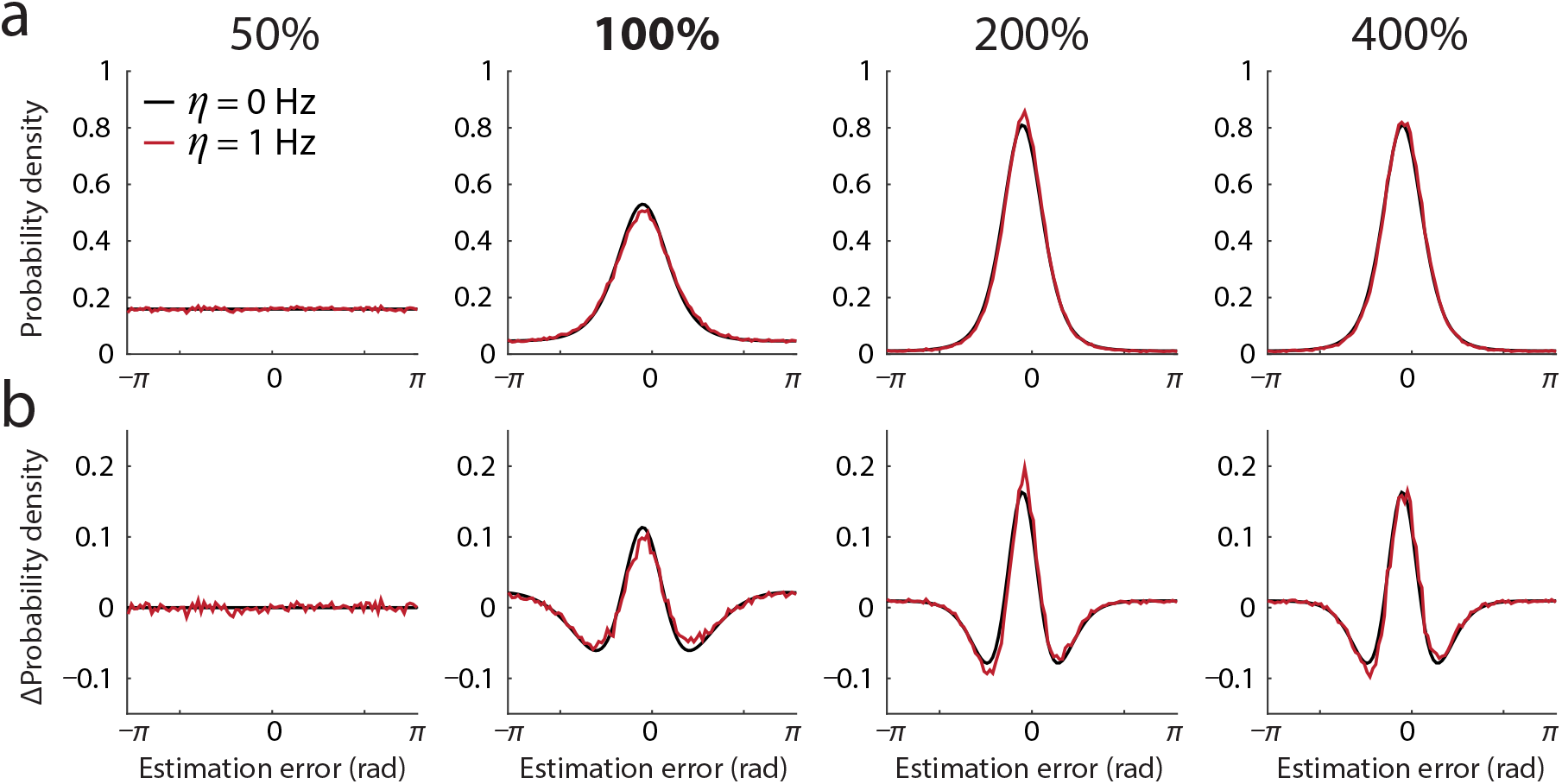
Background (baseline) activity. Black curves show predictions of the population coding model with no baseline activity, for a typical observer. Red curves plot predictions of the model with 1 Hz baseline activity, based on model parameters *γ* and *κ* fit to the no-baseline distributions. Note that with appropriate parameters, the 1 Hz baseline model closely reproduces predictions of the no-baseline model. (a) Error distribution. (b) Deviation from circular normal.

A notable feature of the no-baseline case is the presence of simulated trials in which no spikes occur during the decoding window, and the decoder must ‘guess’ a random value. In the no-baseline model, these trials are prevalent at detection threshold and contribute to the non-normality of the error distribution. In contrast, at threshold, the occurrence of such trials in the baseline case *η* = 1 Hz was negligible (p < 0.0001). This demonstrates that guessing is not critical to generating the non-normal distributions of error observed here, but is rather an artifact of the simplified neuronal model lacking baseline activity.

The model of detection is the same as above, except that now the no-stimulus epoch in general contained spikes, generated at the baseline rate *η*. I used Monte Carlo simulation (discretizing contrast into 100 bins; 10^5^ repetitions, *M* = 100) to estimate the threshold contrast, which again closely approximated the empirical threshold (101% of empirical value for the representative observer). While in the no-baseline case, all errors were due to guesses when no spikes occurred during the stimulus epoch, in the baseline model these trials occurred with negligible frequency (p < 0.0001), providing further evidence that guessing is not a critical element of the population coding model.

## Discussion

The present results demonstrate a signature of population coding in the errors made by human observers in perception of near-threshold stimuli. The predictions of the population coding model reproduce the variability and shape of error distributions in the perceived orientation of a stimulus, as well as capturing the relationship between subjective confidence and perceptual precision. The model also accurately predicted detection threshold based on responses in the reproduction task.

In a recent study (Bays, 2014) I demonstrated deviations from normality, similar to those observed here, in recall errors on a working memory task when memory load was manipulated. The effect of memory set size on precision was explained by a normalization model, in which total population gain was held constant across changes in the number of items represented. Although, as is typical for perceptual tasks, there was a working memory component to the present study, memory limits do not provide an explanation for the present results, as memory load was constant (at one item) across changes in stimulus contrast. Nonetheless, an important alternative hypothesis is that the non-normality observed here arose subsequent to the initial perceptual representation, which itself had normally-distributed error (generated by some unknown mechanism). To test this hypothesis I examined a two-stage model, in which an initial stimulus estimate with normal error was subsequently represented in a population code, with attendant non-normal error. This model failed to reproduce the non-normality in the data, presumably because changes in contrast necessarily had their effect at the initial perceptual stage, where they could not influence the strength of non-normality. This result strongly supports the view that non-normality is present in the initial perceptual representation, and maintained into working memory.

While the population coding model used in the present study incorporates a number of simplifications of the behavior of real neural populations (homogeneity of tuning curves, no baseline activity, no interneuronal correlations), modeling in the working memory study showed that the signature deviations from normality arise independently of these factors. In particular, the model’s behavior was qualitatively unaffected by introducing across-neuron variation in the sharpness of orientation tuning, or changing the shape of the tuning function from Von Mises to cosine. These analyses also identified two factors that serve to increase the population gain corresponding to a given level of variability: the presence of spontaneous (baseline) activity and short-range noise correlations. These factors would prove critical to attaining realistic levels of activity in neural populations on the scale of primary visual cortex. However, analysis of this scenario is hampered by the computational impracticality of simulating activity of hundreds of thousands of correlated neurons.

There is the possibility, in the simplified model of population activity presented here, that no spikes are generated during the decoding interval, resulting in a random response; however, real neurons typically have baseline levels of activity that make this situation unlikely, even at very low contrasts. Additional analysis (Fig 6) confirmed previous modeling work (Bays, 2014) in showing that identical deviations from normality are observed for populations with baseline activity, even though the chance of observing zero spikes is negligible.

The population coding model provided a more parsimonious description of empirical data than a threshold model, in which stimuli are categorically either perceived or not perceived (Luce, 1963; Sergent & Dehaene, 2004; Supèr, Spekreijse, & Lamme, 2001). However, error distributions predicted by the two models were notable mostly for their similarity. Rather than being mutually-exclusive models, I suggest that population coding provides a neural-level explanation for the *appearance* of a threshold in human perception, because the long-tailed error distribution observed at low contrasts resembles a mixture of guessing and accurate judgments.

An interesting outcome of the mathematical analysis presented in the Methods is that the error distributions predicted by the population coding model can be precisely described by an infinite mixture of circular normal distributions. This may provide an explanation for the success of “variable precision” models of working memory (Fougnie, Suchow, & Alvarez, 2012; van den Berg et al., 2012), which attempt to capture recall errors in just such a way, although the proposed distributions over precision in these models do not exactly match that predicted by population coding.

It has long been recognized that our observations are associated with different degrees of certainty, even when the external stimulation that gives rise to the perception is fixed, and further that this certainty is correlated with the magnitude of error in the observation. Clearly we do not have access to the actual error in our observations, or we could correct for it, but exactly what aspect of the perceptual process our sense of confidence is based on is debated (Insabato et al., 2010; Kepecs et al., 2008; Kiani & Shadlen, 2009; Smith & Vickers, 1988). For a population code, an ideal observer of the neural data would base their confidence judgment on the width of the posterior distribution, that is the probability distribution of the stimulus value conditional on the observed spiking activity. I found that this parameter provided an excellent fit to the empirical relationship between subjective confidence and precision of a judgment.

While the posterior width is the best theoretical basis for judging certainty, it is not obvious how it could be computed neurally. The sum of all spiking activity during the decoding window (Bays, 2014; Ma et al., 2006; Pouget, Dayan, & Zemel, 2003) was found to be strongly related both to the width of the posterior and to the precision of the judgment: the more spikes available for decoding, the more precise the estimate. The fit to empirical data was indistinguishable from that using posterior precision, indicating that total spiking activity is a viable and more readily-computable proxy for the true uncertainty in the judgment.

An important limitation of the present study is that both the modeling and experimental work presented here pertain to perception of simple oriented stimuli on a uniform background. Situations in which stimulus energy is present in more than a single orientation, for example encoding an orientation embedded in noise, are not currently represented by the model. Such situations would likely alter the relationship between population activity and precision, potentially making total spike count a less viable basis for subjective confidence. In order to address these issues, future work could expand the encoding model to take arbitrary images as input, perhaps by modeling neural responses as a linear image filtering process followed by a non-linear response transformation (e.g. Goris, Simoncelli, & Movshon, 2015).

## Conclusions

In summary, these results provide behavioral evidence that perception of elementary visual stimuli is an outcome of population coding and decoding at the neural level. Most theoretical work on population codes focuses on the limit of large numbers of spikes, in particular making use of the asymptotic approach to the optimal Cramér-Rao bound (Seung & Sompolinsky, 1993). While some previous studies have analyzed low-spiking regimes (Berens et al., 2011; Brunel & Nadal, 1998; Xie, 2002), they have typically not sought to generate behaviorally-testable predictions. The present results open up the possibility of using analysis of human perceptual reports of near-threshold stimuli to probe the finer details of neural coding that are typically accessible only to non-human electrophysiology. They also have profound implications for signal-detection theory and Bayesian models of perception, which almost universally assume a normal distribution of internal errors.

## Acknowledgments

I thank Peter Dayan for comments on the manuscript, and Leonie Oostwoud-Wijdenes for collecting data. This work was supported by the Wellcome Trust.

